# Neurexin drives *C. elegans* avoidance behavior independently of its post-synaptic binding partner Neuroligin

**DOI:** 10.1101/2024.03.12.584644

**Authors:** Caroline S. Muirhead, Kirthi C. Reddy, Sophia Guerra, Michael Rieger, Michael P. Hart, Jagan Srinivasan, Sreekanth H. Chalasani

## Abstract

Neurexins and their canonical binding partners, neuroligins, are localized to neuronal pre-, and post-synapses, respectively, but less is known about their role in driving behaviors. Here, we use the nematode *C. elegans* to show that neurexin, but not neuroligin, is required for avoiding specific chemorepellents. We find that adults with knockouts of the entire neurexin locus exhibit a strong avoidance deficit in response to glycerol and a weaker defect in response to copper. Notably, the *C. elegans* neurexin (*nrx-1*) locus, like its mammalian homologs, encodes multiple isoforms, α and γ. Using isoform-specific mutations, we find that the γ isoform is selectively required for glycerol avoidance. Next, we used transgenic rescue experiments to show that this isoform functions at least partially in the nervous system. We also confirm that the transgenes are expressed in the neurons and observe protein accumulation in neurites. Furthermore, we tested whether these mutants affect the behavioral responses of juveniles. We find that juveniles (4^th^ larval stages) of mutants knocking out the entire locus or the α-isoforms, but not γ-isoform, are defective in avoiding glycerol. These results suggest that the different neurexin isoforms affect chemosensory avoidance behavior in juveniles and adults, providing a general principle of how isoforms of this conserved gene affect behavior across species.

**Article Summary:** The conserved neurexin locus can encode multiple isoforms via alternate splicing, but very little is known about the function of individual isoforms. We show that the *C. elegans* neurexin γ isoform is specifically required for glycerol, but not copper or quinine avoidance in adults. In contrast, we find that α, but not γ, isoforms are required for avoiding glycerol in juveniles. Collectively, we suggest that different neurexin isoforms are required in juveniles and adults to modify chemosensory behavior.

## Introduction

Animals are exposed to a myriad of chemical cues in their environment. Information about the relevant cues is extracted and processed by the nervous systems, leading to behavioral responses. Studies using insects led to the initial descriptions of repellents and attractants; repellents drive movement away from a chemical source, while attractants drive movements towards the source [1, 2]. Despite rapid progress in identifying the relevant chemical cues, we lack a comprehensive understanding of the neuronal pathways that detect chemical cues and drive behavioral changes. Of particular interest are synaptic proteins, where sensory information is processed to modify animal movement. A complete understanding of this process requires a genetically tractable system with a well-defined nervous system that generates robust behavioral readouts.

The nematode *C. elegans* with just 302 neurons connected by identified chemical and electrical synapses [3], a complete lineage map [4], fully sequenced genome [5], and robust behaviors [6] is ideally suited to reveal how individual genes affect animal behavior. Studies in this model have identified chemosensory receptors [7, 8], neuronal signaling molecules [9], and behavioral models [10-12] required for chemosensory behaviors. Additionally, C. *elegans* has been shown to detect and respond to a wide variety of chemical cues [13, 14]. Notably, this animal has been shown to detect repellents by integrating information from sensory neurons in the head and tail and generate avoidance behaviors [14]. We used repulsion behavior to identify a role for a synaptic protein.

Neurexins and neuroligins are transmembrane proteins that are often localized to neuronal synapses and are thought to be critical components of neuronal circuits [15]. Vertebrates have three neurexin genes (*NRXN1-3*) encoding two isoforms (α, β), and *NRXN1* uniquely encoding a third γ isoform. Also, these genes have multiple splice sites which could result in thousands of unique proteins from the same locus [16]. While these neurexin isoforms have been shown to affect binding partners and may affect synaptic functions [16-18], linking specific isoforms with interacting partners or behaviors has been challenging. In contrast, the *C. elegans* genome encodes a single neurexin gene, *nrx-1*, which encodes α and γ isoforms [19]. Moreover, recent studies suggest that this single locus might also generate several isoforms using alternative splicing [20]. Here we use isoform-specific mutants and show that the γ isoform is selectively required for glycerol avoidance in adults, while the α isoform modifies repellence to the same cue in juveniles.

## Methods

### *C. elegans* strains

All strains were maintained on nematode growth media (NGM) plates with *E. coli* OP50 bacteria as a food source [21]. Larval stage 4 (L4) animals were transferred to new plates with bacteria to maintain each strain. All animals were stored at 20^0^C. A complete list of all strains used in this study and their genotypes are listed in Table S1.

### Avoidance assay

We used a single animal avoidance assay as previously described [14]. Briefly, young adults (day 1 of adulthood) were transferred to a food-free NGM plate and allowed to crawl around for at least 10 minutes before being tested. A small amount of the aversive test compound dissolved in diH_2_O (∼ 5 nL) is placed on the tail of the animal while it is moving forward. Animals that generated a reversal with two or more body bends or a reversal with an omega turn (large-angled turn) upon exposure to the test compound were scored as an avoidance response. Reversals and omega bends have been previously described [22]. A minimum of 6 plates (with 8-15 animals per plates) were tested over three days for each genotype. An avoidance index was calculated as the ratio of number of avoidance responses to the total number of tests.

### Confocal microscopy

*C. elegans* were anesthetized using 5 µL of 100 mM sodium azide in M9 solution, placed on a 5% agarose pad on glass slides, and covered with a glass coverslip. Transgenic *C. elegans* were analyzed using an inverted Leica TCS SP8 laser-scanning microscope operated by LAS X software. Confocal micrographs were generated by compressing z-stacks as maximum intensity projections in FIJI and figures prepared in Adobe Photoshop CS6 and Adobe Illustrator CS6.

### Statistics

Statistical analysis was performed in R using the car and multcomp libraries. The data were fit to a binomial generalized linear model using the glm function. We chose a binomial model because there are two possible behavioral outcomes of exposing a worm to a chemical drop: avoidance or non-avoidance. An ANODEV was subsequently performed on the models using the Anova function. When a significant effect of strain, concentration, or an interaction of both were detected, post hoc analyses were performed using the glht function.

An expected value, predicted avoidance index (PAI), for each strain and condition was calculated using the following equation: log 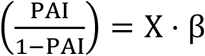. In this equation, X is a design matrix for each strain and concentration. β is a vector of fitted coefficients from the binomial generalized linear model. Since we’re interested in a worm’s response to a stimulus, we can solve the equation for PAI, which will leave us with the following: 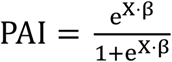. PAI is the avoidance index we expect to find on a given worm strain and concentration. Expected values are included on each graph, depicted as a red line.

## Results

To test the role of neurexin and neuroligin in modifying repellent behaviors, we used a single animal avoidance assay [14]. This assay allows us to identify responses of individual animals to test compounds (**Fig. 1a**, see Methods). We tested two independent alleles in both the *neurexin* and *neuroligin* genes. We found that *neurexin* (*nrx-1*), but not *neuroligin* (*nlg-1*) mutants are defective in their responses to the chemorepellent glycerol. Specifically, we found a deletion of the entire *nrx-1* locus (*wy1155*), or the entire C-terminus of all three isoforms (*wy778*) was deficient in glycerol avoidance (**Fig. 1b**). Notably, an allele deleting about half of the *nlg-1* coding sequence (*ok259*) or one deleting exons 7 and 8 of *nlg-1* (*tm474*) did not significantly alter animal repulsion from glycerol. Next, we tested a *nrx-1, nlg-1* double mutant for glycerol avoidance and found that it was not significantly different from the *nrx-1* glycerol avoidance deficit (**Fig. 1c**). We also detected a significant interaction between strain and glycerol concentration, indicating to us that the strains we tested do not uniformly respond to changes in glycerol concentration. Upon further analysis, we determined that this interaction was due to *nrx-1(*wy778) worms’ decreased response to 1.5M glycerol. To test whether this avoidance deficit was specific to glycerol, we analyzed responses to other widely used chemorepellents, copper and quinine. We saw a slight decrease in copper avoidance in *nrx-1(*wy778) worms while the other neurexin mutant, *nrx-1*(wy1155), trended towards lower avoidance too, this difference however did not survive a multiple corrections analysis (**Fig. 1d)**. We found that all *nrx-1* and *nlg-1* alleles along with the double mutants were indistinguishable from wildtype controls in their responses to quinine (**Fig. 1e**). Taken together, these results show that NRX-1 acts in an NLG-1-independent manner to modify repulsive behavior away from glycerol stimuli, and to a lesser extent, copper. Moreover, this behavioral deficit does not extend to quinine.

**Figure 1:**
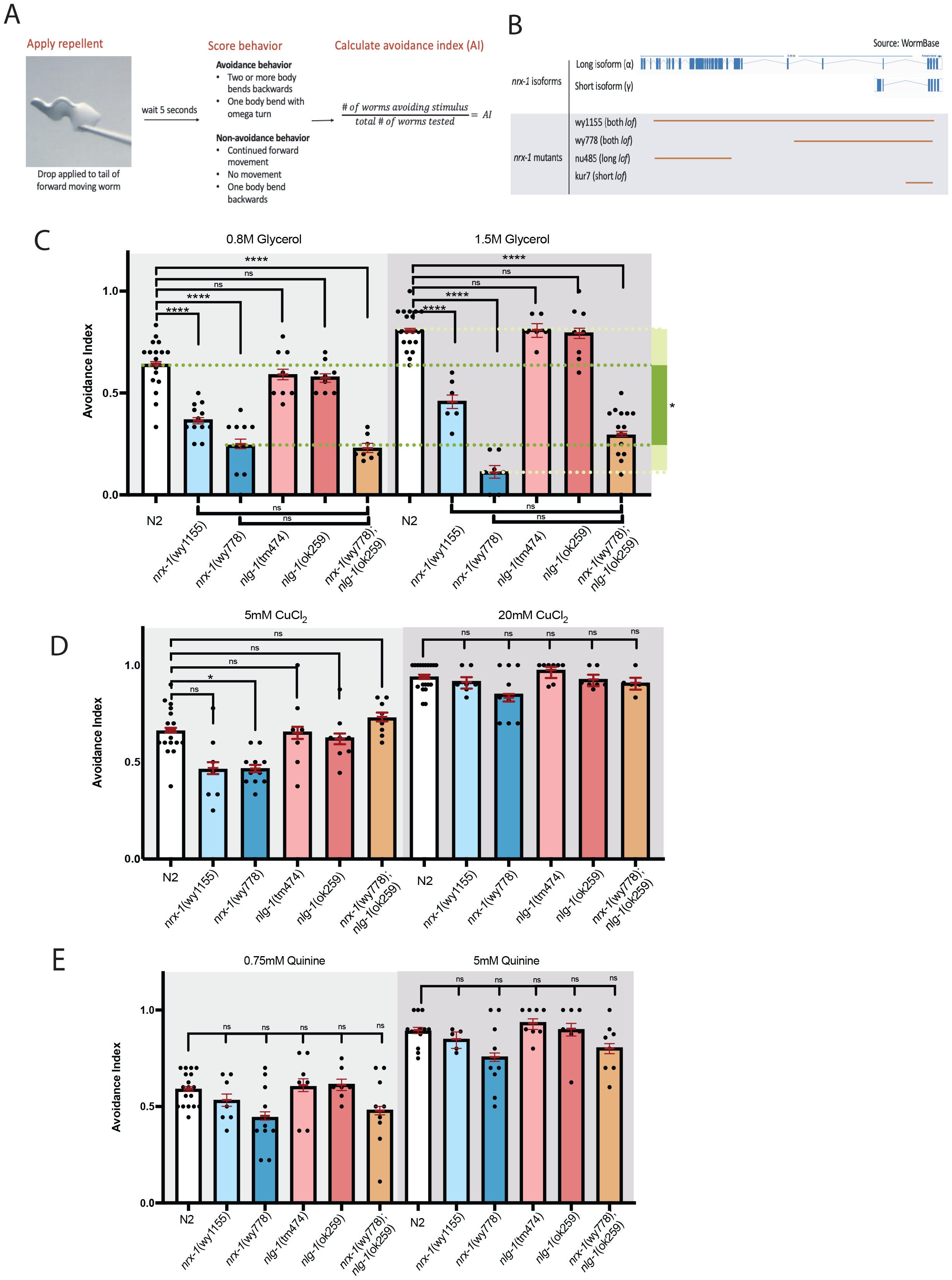
NRX-1 is required for normal glycerol sensation in *C. elegans*. A.) Schematic depicts the workflow for an avoidance assay. To perform the avoidance assay, a drop of repellent is applied to a forward moving worm. The worms’ response is scored for avoidance or non-avoidance. Multiple worms are tested per plate to calculate an avoidance index. B.) Plotted values of avoidance index for each strain in response to 1.5M and 0.8M glycerol. Avoidance index is calculated as number of worms avoiding a drop divided by the number of drops tested per plate. A minimum of six plates was tested per condition. Each black dot represents one plate of 8-15 worms. Error bars are 95% confidence interval. Red lines show predicted/expected avoidance index values based on logistic regression model. We detected significant main effects of strain, concentration, and a significant strain x concentration interaction. Post hoc hypotheses with correction for multiple testing found that *nrx-1*(wy1155) and *nrx-1*(wy778) mutants show a decreased avoidance to 0.8M and 1.5M glycerol. *nlg-1* loss-of-function mutants show no defect. (ns = no significance;*P<0.05, **P<0.01, ***P<0.001, ****P <0.0001). C.) *nrx-1*(wy778) mutants showed a decrease in avoidance to 5mM CuCl_2._ Both *nlg-1* mutants show normal avoidance to 5mM and 20mM copper chloride. We detected significant main effects of strain and concentration but no significant interaction D.) Both *nrx-1* and *nlg-1* mutants show normal avoidance to 0.75mM and 5mM quinine. We detected significant main effects of strain and concentration but no significant interaction.

The *C. elegans nrx-1* locus encodes multiple isoforms, including orthologs of the α and γ isoforms [19, 20]. To test whether specific isoforms affected glycerol avoidance, we analyzed the responses of isoform-specific mutants *nu485*, which knocked out the long α isoform, and *kur7*, which selectively knocks out the γ isoform. We found that *kur7* mutant animals were defective in glycerol avoidance at a lower glycerol concentration (0.8M) (**Fig. 2a**), while *nu485* mutants did not show a significant difference in repulsive behavior at either concentration compared with their wildtype counterparts (**Fig. 2a, 2b**). Next, we tested whether the behavioral deficit in the γ isoform mutants could be rescued by transgenes. We generated transgenic animals expressing the coding sequence for the short γ isoform under a pan-neuronal promoter and analyzed their repulsive responses. We found that neuronal expression of the coding sequence of the γ isoform restored the glycerol avoidance to *kur7* mutants (**Fig. 2a**). We then tested whether the coding sequence of γ isoform can also restore normal behavior to the *wy778* (allele missing the C-terminus of all isoforms) and *wy1155* (allele knocking out the entire *nrx-1* locus). We found that the expression of the short γ isoform in all neurons was able to partially restore normal behavior to both *wy778* (**Fig. 2c**) and *wy1155* alleles (**Fig. 2d**). These results suggest that short γ isoform is selectively required for driving animal avoidance to glycerol. Moreover, this isoform likely functions in the nervous system to affect animal behavior.

**Figure 2:**
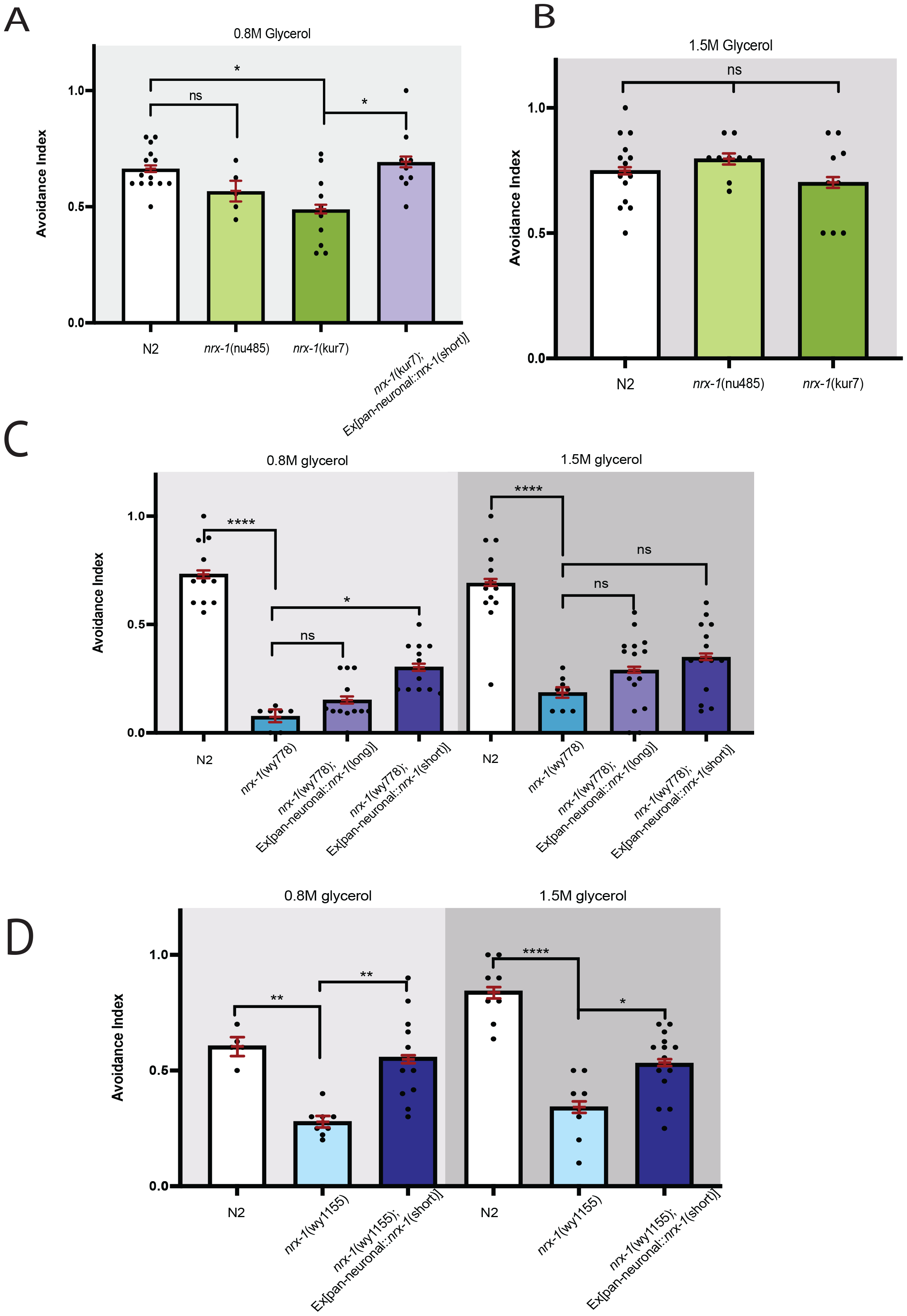
The γ isoform of neurexin primarily modulates glycerol avoidance in adults. A. Plotted values of avoidance index for each strain in response to 0.8M glycerol, as in Figure 1. A minimum of six plates was tested per condition. Each black dot represents one plate of 8-15 worms. Error bars are 95% confidence interval. Red lines show predicted/expected avoidance index values based logistic regression model. We detected a significant effect of strain. The nu485 (α-neurexin *lof* only) shows no avoidance defect to glycerol. The kur7 allele (γ -neurexin *lof* only) shows decreased glycerol avoidance at 0.8M concentration. The defect in the kur7 allele is rescued by expressing β-*nrx-1* under a pan-neuronal promoter. (ns = no significance;*P<0.05, **P<0.01, ***P<0.001, ****P<0.0001). *B. nrx-1*(nu485) and *nrx-1*(kur7) both avoid 1.5M glycerol at the rate of N2. We did not detect a significant effect of strain. Concentrations were analyzed separately because rescue strains were not tested at the higher concentration. C. Expressing α-*nrx*-1 in a *nrx-1*(wy778) background showed no rescue effect. Expressing γ-*nrx*-1 in in a *nrx-1*(wy778) background rescued 0.8M glycerol avoidance. We detected main significant effects of strain and concentration but no significant interaction. D. Expressing γ-*nrx*-1 in in a *nrx-1*(wy1155) background rescued glycerol avoidance at both glycerol concentrations tested. We detected a significant main effect of strain and a significant strain x concentration interaction. Upon post hoc analyses, we saw that both *nrx-1*(wy1155) and its rescue strain showed a trend to not respond to glycerol. The difference between and N2 and *nrx-1*(wy1155) trended towards being different at 0.8M and 1.5M concentrations, however this finding did not survive multiple testing correction.

Our transgenes included a super folded green fluorescent protein (GFP), which allows for the various isoforms to be directly visualized in animals. We analyzed the expression of the γ isoform and compared it with those observed in animals expressing the long α isoform. We found that both fusion proteins were localized to neurites throughout the nerve ring (transgenes were expressed under pan-neuronal promoters) (**Fig. 3**). Additionally, we observed protein accumulation in a punctate fashion in the neurites, with some expression in the cell bodies. These data show that our transgenic proteins are indeed expressed within many if not all neurons in the nervous system.

**Figure 3:**
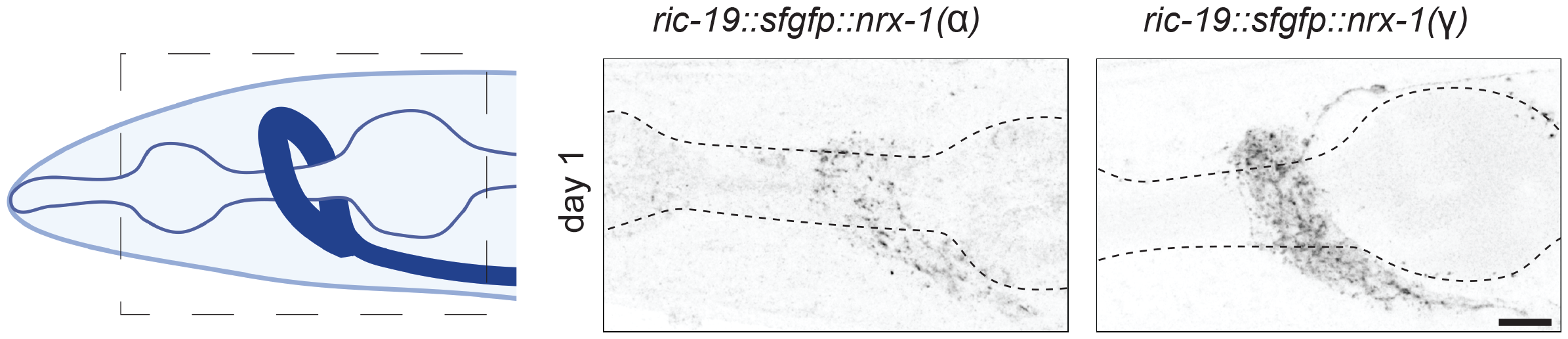
*nrx-1* isoform transgene expression in *C. elegans* neurons. Cartoon of the head of *C. elegans* with pharynx and nerve ring indicated, inset shows location of nerve ring analyzed. Confocal micrograph z-stacks of sfGFP tagged NRX-1 in L3, L4, or adult *nrx-1(wy778)* mutants expressing NRX-1(α) in all neurons or (D) or NRX-1(γ) in all neurons (pseudo-colored black/white, scale bar = 20 micrometers).

Previous studies indicated that neurexins play an important role in assembling synapses [23-25], a critical process in the developing brain. *C. elegans* undergoes a stereotyped developmental program that includes an egg and four larval stages (L1-L4) before molting to an adult [26]. Moreover, each developmental stage includes specific changes to the nervous system including neurogenesis, migration, pruning, and synaptogenesis [27]. To test whether neurexin mutants have phenotypes in developing animals, we first tested whether transgene expression was altered in the developing animal. We observed that expression of both the long α and short γ isoforms were similar to what we observed in the adults (**Fig. 4a**). Next, we analyzed the avoidance responses of alleles that knockout all isoforms, the long α isoform, or the short γ isoform. We found that in both L3 and L4 animals, *nrx-1* alleles knocking out all isoforms (*wy1155* entire locus knockout, and *wy778* knockout deleting C-terminus of all isoforms) were deficient in glycerol response (**Fig. 4b, 4c**). At the L3 stage we saw that both the short and long isoform were individually required for glycerol avoidance (**Fig. 4b**). But interestingly, at the L4 stage the long α isoform *lof* (*nu485*), but not short γ isoform *lof* (*kur7*), were defective in glycerol avoidance (**Fig. 4b, 4c**). These results suggest that different neurexin isoforms may mediate glycerol avoidance in a life stage dependent manner.

**Figure 4:**
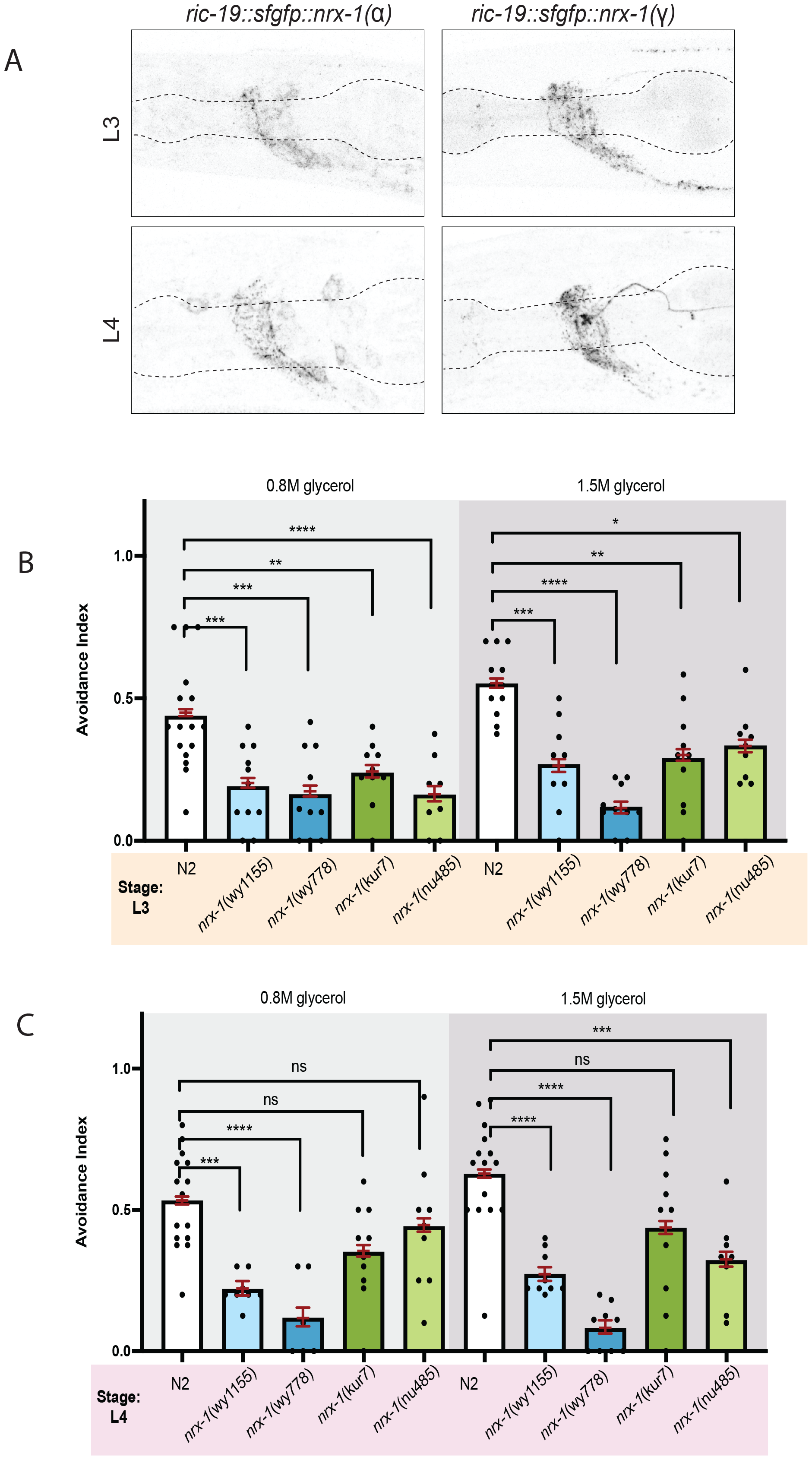
*nrx-1* is required for normal glycerol sensation in L3 and L4 worms. A.) Plotted values of avoidance index for each strain in response to 0.8M and 1.5M glycerol, as in Figure 1. A minimum of six plates was tested per condition. Each black dot represents one plate of 8-15 worms. Error bars are 95% confidence interval. Red lines show predicted/expected avoidance index values based on binomial linear model. We detected main effects of strain and concentration but no significant interaction. *nrx-1* wy1155, wy778, kur7, and nu485 L3 worms show reduced glycerol avoidance compared to N2. (ns = no significance;*P<0.05, **P<0.01, ***P<0.001, ****P <0.0001) *B*.) At the L4 stage, wy1155 and wy778 showed reduced glycerol avoidance at both concentrations. The kur7 allele showed no defect while the nu485 allele showed a defect in avoidance at 1.5M glycerol. We detected a main effect of strain.

## Discussion

Our study shows that the short neurexin γ isoform drives glycerol avoidance in adults, while long α neurexin isoform play a larger role in glycerol avoidance in juvenile worms. Moreover, avoidance deficits in neurexin mutants are the strongest in response to glycerol but there is also a slight defect in copper sensation. Previous studies using cell ablations and calcium imaging have revealed the sensory neurons that are critical to detecting these repellents. While ASH sensory neurons are required for detecting glycerol [28], ASH and ASK neurons are required for quinine [29], and ASH, ADL, and ASE neurons are required for copper avoidance [30]. We suggest that the γ isoform might be selectively required in ASH neurons and predict that animals missing this isoform have defective ASH-driven avoidance behaviors, consistent with our observed glycerol avoidance deficit. However, since quinine and copper can also be detected by additional sensory neurons, we do not observe strong avoidance deficits to those cues.

Our results also show that NRX-1 functions in an NLG-1-independent manner to modify glycerol avoidance. These data are somewhat surprising since both of these proteins are thought to bind each other [24] and function together [31, 32]. However, additional studies have shown that these proteins can function independently [33-35], or even antagonistically [36, 37]. Our results are consistent with the growing consensus that NRX-1 can have functions that are independent of NLG-1. We speculate that the short γ NRX-1 isoform might interact with other binding partners, including calystenins (*casy-1*), dystroglycans (*dgn-1/2/3*), GluK/GRIK (*glr-1/2/3/5*), GluD1 (*gdh-1*), MDGA1 (*igcm-1*), GABRA1 (*lgc-37*), LRRTM1/2 (*lron-3, sma-10, hmp-1, pan-1*), and others [31, 33, 35, 37], whose homologs are encoded in the *C. elegans* genome.

We also identify a differential role for the neurexin isoforms during development. Specifically, we find that the long α isoform is required for glycerol avoidance in L4 worms, while the short γ isoform is required for avoiding this cue as an adult. We previously showed that *C. elegans* sensory circuits undergo a juvenile-to-adult transition. Specifically, it was found that while ASH neurons were required to avoid high concentrations of diacetyl in adults, ASH, AWA, and ASK were required to avoid this same cue in juvenile L3 animals [38]. Consistent with this data, we suggest that neural circuit detecting glycerol undergoes a juvenile-to-adult transition and speculate that the long α isoforms is selectively required in the juvenile neural circuit for glycerol repulsion. More broadly, we suggest that the developing adolescent and mature adult brains may utilize different neurexin isoforms to drive behavior.

## Acknowledgements

We thank P. Kurshan for the γ isoform specific mutant and CGC for other strains. The authors also thank members of the Chalasani, Srinivasan, and Hart labs for technical assistance and feedback on this project.

## Funding

This work was supported in part by grants from NIH 1R35GM146782 (MPH), 1R01DC016058 (JS), 1R01MH096881 (SHC), and Nippert Foundation (SHC). Some strains were provided by the Caenorhabditis Genetics Center, which is funded by NIH Office of Research Infrastructure Programs (P40 OD010440).

## Author Contributions

CSM, JS, and SHC conceived and designed the study and experiments, and CSM and SG conducted all behavioral experiments. KCR generated transgenic animals and performed genetic studies. MPH designed experiments and generated data for the localization study. MR provided statistical analysis guidance and baseline R code. CSM wrote the manuscript with assistance from SHC and JS, and all authors, reviewed, revised, and approved the manuscript.

## Conflict of Interest

The authors declare no conflicts of interest.

## Figure legends

**Supplementary Table S1**. Table showing the list of strains used in this study along with their genotypes.

**Table 1:** List of strains used this study along with their genotypes.

| Strain name | Genotype |
| --- | --- |
| TV13570 | *unc-119(ed3) III; nrx-1(wy778[unc-119(+)]) V* |
| TV22998 | *nrx-1(wy1155) V* |
| VC228 | *nlg-1(ok259) X* |
| IV91 | *nlg-1(tm474) X* |
| IV936 | *nrx-1(wy778[unc-119(+)]) V; nlg-1(ok259) X* |
| TV22997 | *nrx-1(nu485) V* |
| PTK69 | *nrx-1(kur7) V* |
| IV891 | *nrx-1(wy778) V; ueEx617[ric-19p::nrx-1; unc-122p::RFP]* |
| MPH15 | *nrx-1(wy778) V; hpmEx4[ric-19p::nrx-1(short); myo-2::rfp]* |
| IV1043 | *nrx-1(wy1155) V; hpmEx3[ric-19p::nrx-1(m-short); myo-2::rfp]* |
| IV1066 | *nrx-1(kur7) V; hpmEx4[ric-19p::sfGFP::nrx-1(m-short); myo-2::rfp]* |

